# Patterns of lineage-specific genome evolution in the brood parasitic black-headed duck (*Heteronetta atricapilla*)

**DOI:** 10.1101/2022.05.22.492970

**Authors:** Sara J. Smith, LaDeana W. Hillier, Christopher N. Balakrishnan, Michael D. Sorenson, Wesley C. Warren, Timothy B. Sackton

**Affiliations:** Informatics Group, Faculty of Arts and Sciences, Harvard University, Cambridge, MA, 02138, USA; Department of Biology, Boston University, Boston, MA, 02215, USA; Department of Animal Sciences, Institute for Data Science and Informatics, Bond Life Sciences Center, University of Missouri, Columbia, MO, 65211, USA; Department of Biology, East Carolina University, Greenville, NC, 27858, USA

## Abstract

Occurring independently at seven separate origins across the avian tree of life, obligate brood parasitism is a unique suite of traits observed in only approximately 1% of all bird species. Obligate brood parasites exhibit varied physiological, morphological, and behavioural traits across lineages, but common among all obligate brood parasites is that the females lay their eggs in the nest of other species. Unique among these species is the black-headed duck (*Heteronetta atricapilla*), a generalist brood parasite that is the only obligate brood parasite among waterfowl. This provides an opportunity to assess evolutionary changes in traits associated with brood parasitism, notably the loss of parental care behaviours, with an unspecialized brood parasite. We generated new high-quality genome assemblies and genome annotations of the black-headed duck and three related non-parasitic species (freckled duck, African pygmy-goose, and ruddy duck). With these assemblies and existing public genome assemblies, we produced a whole genome alignment across Galloanserae to identify conserved non-coding regions with atypical accelerations in the black-headed duck and coding genes with evidence of positive selection, as well as to resolve uncertainties in the duck phylogeny. To complement these data, we sequenced a population sample of black-headed ducks, allowing us to conduct McDonald-Kreitman tests of lineage-specific selection. We resolve the existing polytomy between our focal taxa with concordance from coding and non-coding sequences, and we observe stronger signals of evolution in non-coding regions of the genome than in coding regions. Collectively, the new high-quality genomes, comparative genome alignment, and population genomics provide a detailed picture of genome evolution in the only brood parasitic duck.

## Introduction

About 1% of bird species are obligate brood parasites (Rothstein 1990), which reproduce only by exploiting the parental care behaviour of other species (Roldán & Soler 2011; Soler 2017). While this social parasitism is accomplished via the seemingly simple behaviour of depositing eggs in the nests of an appropriate host species, the ensuing interactions between hosts and parasites have produced a diversity of fascinating adaptations and counter-adaptations (Stoddard & Hauber 2017; Feeney et al. 2012). Shared in common among all obligate brood parasites is the convergent loss of all parental care behaviour, including nest building, incubation and provisioning/guarding offspring, all of which represent fundamental, ancestral behaviours in most birds (Soler 2017; Davies 2010). Given seven independent origins of obligate brood parasitism in birds (Sorenson & Payne 2002), there is an excellent opportunity to explore the genomic correlates and consequences of this major behavioural and life history transition. We focus here on a genomic comparison of the brood parasitic black-headed duck *Heteronetta atricapilla* with representatives of three closely related waterfowl genera with typical parental care behaviours.

The black-headed duck is a generalist brood parasite that lays eggs in the nests of a wide-range of marsh-nesting host species, but relies primarily on two species of coots (genus *Fulica*) to rear its offspring (Lyon & Eadie 1991; Cabrera et al. 2017). It is the only obligate brood parasite among the waterfowl (Anseriformes), as well as the only avian obligate brood parasite with precocial offspring. Indeed, black-headed duck ducklings require minimal post-hatch parental care, typically leaving the host nest within a day of hatching and thereafter fending for themselves (Weller 1968; Lyon & Eadie 2013; Rees & Hillgarth 1984). This is in marked contrast to the altricial offspring of other avian brood parasites. Black-headed ducks do not exhibit obvious host-specific adaptations, such as mimicry of host egg coloration as is observed in many other obligate brood parasites (Davies 2010; Stoddard & Hauber 2017). Interestingly, facultative intraspecific brood parasitism in the two primary hosts (*Fulica armillata* and *F. rufifrons*) and the associated evolution of their egg recognition capabilities may have precluded black-headed ducks from evolving egg mimicry (Lyon & Eadie 2004).

While the black-headed duck is unique among waterfowl in having lost parental care behaviour, it remains phenotypically similar to other duck species. Past research, however, has focussed on its behavioural ecology and little is known about possible physiological or developmental adaptations that might be associated with the evolution of obligate parasitism. Unlike some other brood parasites (Igic et al. 2011; López et al. 2018), there is no evidence that *H. atricapilla* has evolved thicker eggshells (Mallory 2000). Other possible axes of evolutionary change relevant to all obligate brood parasites include relaxed selection associated with the loss of parental care behaviors (Lynch et al. 2019) and adaptations for life as a parasite, such as increased fecundity (Payne 1974; Scott & Ankney 1983) and enhanced spatial abilities associated with finding and tracking the status of host nests (Sherry & Guigueno 2019). For black-headed duck in particular, there may be physiological or developmental changes associated with what appears to be enhanced precociality in the parasitic ducklings.

Here, we compare genome evolution in black-headed duck with three species representing the most closely related waterfowl genera with typical nesting and parental care behaviour to test whether black-headed duck is an outlier with respect to various measures of genome evolution. We test for accelerated evolution in both protein-coding sequences and conserved non-exonic elements (CNEEs), and ask whether such changes in black-headed duck are overrepresented in particular genetic pathways or gene ontology (GO) categories.

## Methods

### Genome assembly and whole genome alignment

Blood samples were taken from female individuals of black-headed duck *Heteronetta atricapilla*, freckled duck *Stictonetta naevosa*, ruddy duck *Oxyura jamaicensis*, and African pygmy goose *Nettapus auritus* at the Sylvan Heights Bird Park (Scotland Neck, North Carolina). Using the Supernova pipeline (Weisenfeld et al. 2017), the Illumina BCL output was used to generate FASTQs from which genomes were assembled *de novo* with pseudohaplotypes written as separate FASTAs. Chromosomes were ordered and oriented using mallard *Anas platyrhynchos* and chicken *Gallus gallus* assemblies as guides (Table S2). Our genome assemblies for the four duck species are publicly available on NCBI (BioProject PRJNA588796).

To facilitate downstream comparative analyses, we used Progressive Cactus (Armstrong et al. 2020) to generate a whole genome alignment for our focal taxa and eleven additional representatives of Galloanserae (Table S2). This alignment included nine anatid species (including our four focal taxa) and six galliformes, representing the full complement of currently available (as of January 2020), high quality Galloanserae genome assemblies. The quality of our genome assemblies (Table S1) was assessed by Benchmarking Universal Single Copy Orthologs (BUSCO) against the Aves database (database v10, Seppey et al. 2019).

### Genome annotation

To generate genome annotations using Comparative Augustus (Stanke et al. 2008), we used publicly available RNA-seq data for *O. jamaicensis* (Schneider et al. 2019), NCBI BioProject PRJNA517454) to generate a splice-aware gapped alignment using a two-pass iterative mapping approach. This alignment along with existing genome annotations for *G. gallus, A. platyrhynchus*, and *O. jamaicensis* (the new genome we report here was annotated by NCBI using the same RNA-seq data noted above) were integrated into an SQL hints database. The whole genome alignment was split into sequence blocks to allow for parallel processing of gene predictions. Gene predictions from parallel runs of Comparative Augustus CPG (König et al. 2016) were merged to produce the final annotations. Genome annotation completeness (Table S1) was estimated by Benchmarking Universal Single Copy Orthologs (BUSCO) against the Aves database (database v10, Seppey et al. 2019).

### Variant calling

To support population genomic analyses, we generated resequencing data for fourteen black-headed ducks (12 females and two males) originating from Buenos Aires Province, Argentina (see Lyon & Eadie 2013 for more detailed locality information). Black-headed duck DNA extracts were provided by colleagues at University of California Davis. We also generated resequencing data for one male of each of the three outgroup species, using blood samples from captive birds at Sylvan Heights Bird Park. Fragment library preparation and sequencing (DNBSEQ) were completed by BGI Genomics; 100 base-pair, paired-end reads were generated for each sample at 10-15x coverage for the 12 female samples and at 33-38x coverage for the five male samples. Sequencing data from the four females used for genome assembly, the resequencing data for males of the three nesting species, and the population sample of black-headed ducks were processed with *snpArcher* (https://github.com/harvardinformatics/snpArcher), a reproducible pipeline that maps reads to the reference genomes and calls variants with GATK (Van der Auwera & O’Connor 2020), resulting in species-specific Variant Call Format (VCF) files with GATK hard filtering annotations, coverage calculations, and missingness statistics. The resequencing data from the male *S. naevosa* was also aligned to the black-headed duck genome using *snpArcher* for downstream analyses requiring an outgroup taxon.

### Phylogenetic relationships

Traditionally regarded as a member of the waterfowl tribe Oxyurini (Livezey 1995), the phylogenetic relationships and closest nesting relative(s) of the black-headed duck remain uncertain because no published analysis of molecular data has included all the relevant taxa needed for a robust test (Gonzalez et al. 2009; Sun et al. 2017). Our choice of outgroup taxa for genome assembly was based on unpublished analyses (Sorenson et al., unpubl. data) suggesting monophyly of a clade comprising the four genera represented by our focal taxa plus *Nomonyx*, the well-supported sister genus to *Oxyura* (e.g., Gonzalez et al. 2009; Sun et al. 2017). To further test this clade and to identify the black-headed duck’s closest living relative, we constructed and analysed three data sets:

1. We assembled complete mitochondrial DNA (mtDNA) sequences for one individual of each of our four focal species and combined these with 29 additional anseriform mt genomes from GenBank, including magpie goose *Anseranas semipalmata* and southern screamer *Chauna torquata* to root the tree (see Table SX for a full list of taxa and GenBank accession numbers). The data were partitioned by gene and codon position and analysed in PartitionFinder 2 (Lanfear et al. 2016) to develop an appropriate partitioning strategy. The data were then analysed in BEAST v. 2.4.6 (Bouckaert et al. 2019).
2. For tree estimation with non-coding sequences, the full set of CNEEs were filtered for elements over 500 bp and those that are present in all Galloanserae species from the whole genome alignment. The filtered set of CNEEs were aligned with MUSCLE (Edgar 2004) and alignments were trimmed manually in Geneious (Kearse et al. 2012) to remove sites where indels were present in more than one of the species. The aligned CNEEs were analysed with RAxML-NG (Kozlov et al. 2019) to infer the best tree for each sequence, both with and without bootstrapping. The consensus species tree was estimated in ASTRAL (Zhang et al. 2018) from the unrooted gene trees.
3. To estimate the phylogenetic relationship with coding sequences, we used protein-translated annotations as well as nucleotide sequences of coding regions. We translated the publicly available and Comparative Augustus genome annotations for the Galloanserae species to amino acid sequence with *gffread* (Pertea & Pertea 2020) and estimated the species tree with OrthoFinder (Emms & Kelly 2019). We corroborated the OrthoFinder species tree by independently analysing the gene trees from OrthoFinder with ASTRAL (Zhang et al. 2018). To prepare the nucleotide sequences of coding regions, we translated the genome annotations from Comparative Augustus (Stanke et al. 2008) to chicken-reference gene symbols using the OrthoFinder (Emms & Kelly 2019) orthologue translation files. Nucleotide sequences for coding regions were extracted as gene-specific FASTAs, aligned with MAFFT (Katoh & Standley 2013), and cleaned with the segment filtering software HmmCleaner (Di Franco et al. 2019) before analysing with RAxML-NG (Kozlov et al. 2019) and ASTRAL (Zhang et al. 2018) as described above for the consensus tree estimation in non-coding sequences.

### Signatures of selection in non-coding sequences

Since regulatory regions are known to underlie some examples of phenotypic evolution among species (Sackton et al. 2019, e.g., obligate brood parasitism), we built a set of consensus conserved non-exonic elements (CNEEs) from published sets of vertebrate CNEEs (Craig et al. 2018; Lowe et al. 2014; Sackton et al. 2019; Siepel et al. 2005). The individual sets of CNEEs were lifted over to *G*.*gallus* (GRCg6a) coordinates using halLiftover (Hickey et al. 2013) and the assembled consensus sequences were aligned with MAFFT (Katoh & Standley 2013). The aligned sequences were concatenated with CatSequences (doi: 10.5281/zenodo.4409153) and 4-fold degenerate neutral models were generated with PHAST (Hubisz et al. 2011). The total set of consensus CNEEs were assessed for evidence of conservation using phyloP (Hubisz et al. 2011) and any CNEEs that did not show evidence of conservation were removed from the consensus set. With the remaining CNEEs (over 375,000 elements), we used PhyloAcc (Hu et al. 2019) to identify evolutionary rate accelerations in each focal species.

The set of accelerated CNEEs in each focal species were analysed to identify i) spatial clusters of accelerated CNEEs across the genome, ii) genes enriched for accelerated CNEEs in nearby regions, and iii) Gene Ontology (GO) terms enriched for an overrepresentation of accelerated CNEEs. To identify spatial enrichment of accelerated CNEEs, all CNEEs were binned in 100 kb windows across the genome with a 50 kb sliding window and a used a binomial test to calculate whether there is an excess of accelerated CNEEs relative to the genomic background in the window. To assess the enrichment of accelerated CNEEs in nearby regions of genes and the overrepresentations of GO terms, permutation tests were conducted on 100 kb windows by comparing the observed data with null background distributions (Yu et al. 2012; Durinck et al. 2009).

### Signatures of selection in coding sequences

Orthogroups that contained *H. atricapilla*, were represented in at least 50% of the Galloanserae species from the whole-genome alignment, and contained less than 25 sequences were selected to test for signatures of selection in coding regions. These protein alignments from OrthoFinder were translated to nucleotide alignments using the genome annotations produced by either Comparative Augustus or NCBI. The nucleotide alignments were then corrected for frame-shift mutations and translated back to protein sequences using the codon-aware multiple sequence alignment (MSA) pre-processing script (described in Kosakovsky Pond & Frost 2005), aligned with MUSCLE (Edgar 2004), then mapped back to nucleotide sequences using the codon-aware MSA post-processing script (Kosakovsky Pond & Frost 2005) without compressing duplicate sequences. These MSAs were then trimmed for duplicate sequences, using the gene tree for each orthogroup and then cleaned with the segment filtering software HmmCleaner (Di Franco et al. 2019). Gene trees for all MSAs were pruned using *Newick Utils* (Junier & Zdobnov 2010), which were used as guide trees in running each alignment through an adaptive branch-site random effects likelihood analysis (aBSREL, Smith et al. 2015) and a branch-site unrestricted statistical test for episodic diversification analysis (BUSTED, Murrell et al. 2015) from the Hypothesis Testing using Phylogenies (HyPhy) software suite (Pond et al. 2004).

With the population resequencing data for *H. atricapilla* and the genome sample of *S. naevosa*, we used a custom pipeline to conduct McDonald-Kreitman (MK) tests for signatures of selection in genes (McDonald & Kreitman 1991). We translated the *H. atricapilla* coding regions from the Comparative Augustus genome annotation to the corresponding *G. gallus* gene names and built a functional annotation database in snpEff (Cingolani et al. 2012). In brief, the pipeline filtered the VCFs, recalculated coverage, created a set of callable sites common between the ingroup (*H. atricapilla*) and the outgroup (*S. naevosa*), functionally annotated the variants in the ingroup and the outgroup using the custom snpEff database, parsed the synonymous and nonsynonymous variants, associated those variants with genes, built *d*_*n*_*/d*_*s*_ tables for each gene, and conducted the traditional MK test, an extension of the MK test in SnIPRE (Eilertson et al. 2012), as well as calculated the population statistic *alpha* (the proportion of amino acid substitutions fixed by positive selection, Eyre-Walker 2006; McDonald & Kreitman 1991) and the direction of selection calculation for each gene (Stoletzki & Eyre-Walker 2011).

To assess if there is an enrichment of GO terms associated with obligate brood parasitism, we used both a foreground/background approach as well as a gene set enrichment (GSE) analysis. For the GSE approach, we used both the *G. gallus* database, as well as lifting over the coding regions to human coding regions for use with the *Homo sapiens* database.

To better understand the molecular evolution of *H. atricapilla*, we also inferred demographic history and scanned for selective sweeps. To estimate the demographic history from sequence data, we calculated the folded site frequency spectrum using sites that were present in all individuals. Using an estimated mutation rate per site per generation from the collared flycatcher (*Ficedula albicollis*, Smeds et al. 2016) and the number of callable sites in the genome, we used Stairbuilder to generate a Stairway plot (Liu & Fu 2020) of *H. atricapilla*. In order to scan the genome for selective sweeps using SweepFinder2 (DeGiorgio et al. 2016), we generated allele frequency files for each major chromosome, as well as the genome-wide empirical frequency spectrum. Scans were run on each major chromosome in 1 kb windows.

All the annotated code for the analyses described above are publicly available on GitHub at https://github.com/sjswuitchik/duck_comp_gen.

## Results

### Genome assembly and annotation

All four focal taxa genomes were assembled at 41.7x to 54.96x raw coverage with a contig N50 between 132.48 Kb and 155.79 Kb, resulting in assemblies between 1.06 Gb and 1.1 Gb (Table S1). Over 95% of each genome is assembled in scaffolds larger than 10 Kb with an estimated BUSCO completeness score of 94.6-95% (Table S1). The genome annotations produced by Comparative Augustus CPG are estimated to be 90.6% to 92.6% complete (Table S1).

### Phylogenetic polytomy resolution

The OrthoFinder species tree place *Heteronetta atricapilla* sister to *Stictonetta naevosa*, with *Oxyura jamaicensis* sister to this clade and *Nettapus auritus* sister to the ducks. However, the proportion of gene trees supporting the relationships were quite low (0.195 - 0.251 of 17,184 trees). When analysing the gene trees which contained only single copy orthologs (*n =* 5682) with RAxML-NG and drawing a consensus tree with ASTRAL, this relationship is supported with local posterior probabilities of 1 for each node (Fig. 1). This relationship was also seen in analysing protein coding alignments (*n =* 12,133) with RAxML-NG and a consensus tree drawn with ASTRAL. This relationship was further supported by the parsimony analysis of 6000 CNEEs that were present in all species of the tree, where all nodes have 100% bootstrap support except for the *H. atricapilla-S*.*naevosa* node, where the bootstrap support is 99%. The RAxML-NG analysis of the same CNEEs and ASTRAL consensus tree also supported this relationship with local posterior probabilities of 1 for each node. The BEAST analysis of the mitogenomes did not resolve the polytomy, but did place *N. auritus* outside of the trichotomy of the remaining focal taxa with a posterior probability of 1.

**Figure 1.**
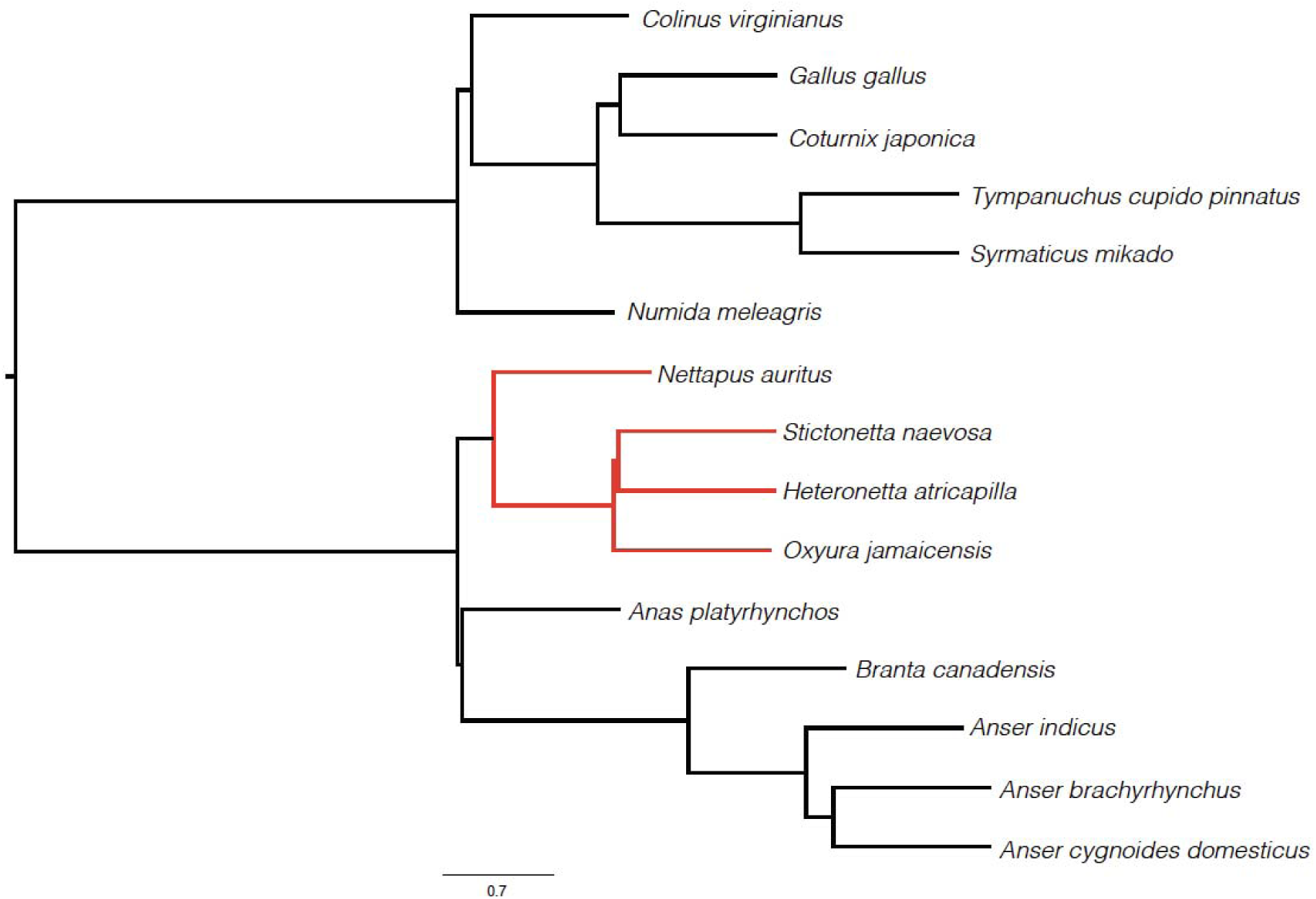
Consensus phylogenetic tree of fifteen Galloanserae species from publicly available data, including the four focal Anseriformes genomes, resolved with both protein-coding and conserved non-exonic element sequences.

### Signatures of selection in non-coding sequences

There were a similar number of accelerated CNEEs identified in each focal species (Table SX), with slightly more in both *H. atricapilla* and *N. auritus*. In scanning the genome for spatial enrichment of accelerated CNEEs, there were no large, concentrated clusters identified in any of the focal taxa. There are some windows of interest in *H. atricapilla* (Fig. 2A), *S. naevosa*, and *N. auritus* on the Z chromosome (Fig. S1A,C) but there were no windows with a significant number of accelerated CNEEs in *O. jamaicensis* (Fig. S1B).

**Figure 2.**
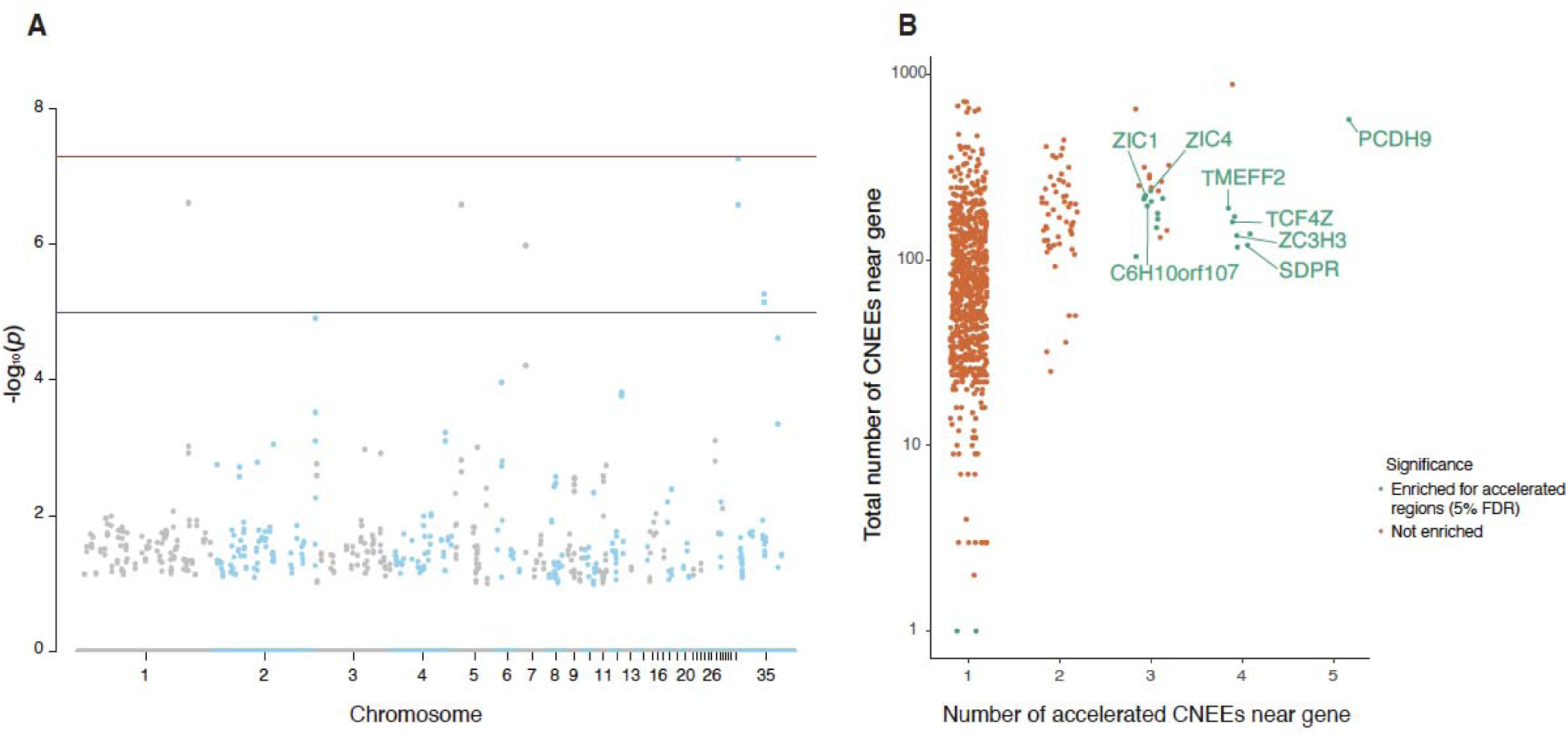
Signatures of selection in the black-headed duck. A) Genome-wide scan for clusters of accelerated conserved non-exonic elements (CNEEs) and B) genes significantly enriched for accelerated regions of CNEEs.

There are 21 genes that are enriched for accelerated regions in *H. atricapilla* (Fig. 2B), which are primarily involved in neurogenesis and neuronal development (5/21), and calcium-dependent processes (2/21). The genes involved in neurogenesis that are enriched for accelerated regions include Zic Family Members 1 and 4 (ZIC1, ZIC4), both of which are associated with the cerebellum development; *transcription factor 4* (TCF4), which is involved in the initiation of neuronal differentiation; and *protocadherin 9* (PCDH9), a protein which mediates cell adhesion in neural tissues. These transcription factors with roles in brain development are enriched for accelerated regions specifically in *H. atricapilla*, and not in the other related nesting species.

However, the majority of these 21 enriched genes (13/21) are uncharacterized loci without functional annotations. There are no genes enriched for accelerated regions in *S. naevosa* and *O. jamaicensis* (Fig. S2A,B) however, there are 126 enriched genes in *N. auritus* (Fig. S2C) that are associated with neuronal development (12/162), photoreception (8/162), and immunity (5/126).

In analysing the overrepresentation of GO terms associated with the 294 accelerated CNEEs in *H. atricapilla*, there are 21 significantly enriched GO terms across all three subontologies (Biological Process, BP; Cellular Component, CC; and Molecular Function, MF; Table 1). The most prominent signal in these terms is related to olfaction, including sensory perception of smell (GO:0007608, BP, p = 0.036), odorant binding (GO:0005549, MF, p = 0.035), and olfactory receptor activity (GO:0004984, MF, p = 0.035). These terms are all functionally associated with *olfactory receptor 6* (OLFR6), an olfactory receptor gene on chromosome 5.

**Table 1.** Gene Ontology (GO) terms associated with enriched CNEEs in black-headed duck.

| GO ID | GO Annotation | Subontology | p |
| --- | --- | --- | --- |
| GO:0007155 | cell adhesion | BP | 0.00799201 |
| GO:0009887 | animal organ morphogenesis | BP | 0.00799201 |
| GO:0022610 | biological adhesion | BP | 0.00799201 |
| GO:1901564 | organonitrogen compound metabolic process | BP | 0.01298701 |
| GO:0019538 | protein metabolic process | BP | 0.01898102 |
| GO:0034329 | cell junction assembly | BP | 0.02897103 |
| GO:0051301 | cell division | BP | 0.03196803 |
| GO:0007608 | sensory perception of smell | BP | 0.03596404 |
| GO:0050911 | detection of chemical stimulus involved in<br>sensory perception of smell | BP | 0.03596404 |
| GO:0043604 | amide biosynthetic process | BP | 0.03796204 |
| GO:0055114 | oxidation-reduction process | BP | 0.03996004 |
| GO:0050907 | detection of chemical stimulus involved in<br>sensory perception | BP | 0.04095904 |
| GO:0044267 | cellular protein metabolic process | BP | 0.04695305 |
| GO:0044248 | cellular catabolic process | BP | 0.04895105 |
| GO:0005794 | Golgi apparatus | CC | 0.00899101 |
| GO:0016020 | membrane | CC | 0.03096903 |
| GO:0071944 | cell periphery | CC | 0.04495505 |
| GO:0070161 | anchoring junction | CC | 0.04595405 |
| GO:0140096 | catalytic activity, acting on a protein | MF | 0.02597403 |
| GO:0004984 | olfactory receptor activity | MF | 0.03496504 |
| GO:0005549 | odorant binding | MF | 0.03496504 |

There are 19 significantly enriched GO terms in *S. naevosa* and 21 significantly enriched GO terms in *O. jamaicensis* (Table S3). The GO terms enriched in both of these focal taxa are fairly general, however there are 3 GO terms associated with hemoglobin complexes found in *O. jamaicensis*. In the most divergent focal taxa with the longest branch length, *N. auritus*, there are 195 significantly enriched GO terms, predominantly in the BP ontology (*n*_*BP*_ = 102, *n*_*CC*_ = 15, *n*_*MF*_ = 78, Table S3). The most prominent annotations are related to immunity (e.g., GO:0043123, positive regulation of NF-kappaB signalling, p = 0.032; GO:0002755, myD88-dependent toll-like receptor signalling pathway, p = 0.018; GO:0150076, neuroinflammatory response, p = 0.023), kidney development (e.g., GO:0072163 and GO:0072164, mesonephritic epithelium and tubule development, respectively, p = 0.041), and regulation of nervous system development (e.g., GO:0050768, negative regulation of neurogenesis, p = 0.028; GO:0051961, inhibition of nervous system development, p = 0.029; GO:0045665, negative regulation of neuron differentiation, p = 0.044).

### Signatures of selection in coding sequences

From the HyPhy analyses, we defined an orthogroup as being of interest if i) aBSREL identified significant episodic selection on the *H. atricapilla* branch (p < 0.05 after FDR correction), ii) BUSTED also identified positive selection in this orthogroup (p < 0.05 after FDR correction), and iii) there were not more than two species in addition to *H. atricapilla* in the phylogeny that were identified to have experienced significant episodic selection by aBSREL. There were 151 orthogroups that fit the above set of criteria but in analysing the overrepresentation of GO terms associated with these orthogroups of interest, there were no significantly enriched GO terms across all subontologies. There was also no overlap between two lists of *a priori* parental care genes (Hackett et al. 2008; Lynch et al. 2019) and the orthogroups of interest.

From polymorphism and divergence data calculated per gene in *H. atricapilla* with *S. naevosa* acting as the outgroup, the direction of selection (DoS) was calculated for each gene as the direction and degree of departure from expectations of neutral evolution (Stoletzki & Eyre-Walker 2011). The distribution of the DoS statistic suggests predominantly neutral and negative selection (Fig. 3A). The distribution of *alpha* per gene, the proportion of amino acid substitutions fixed by positive selection (McDonald & Kreitman 1991; Eyre-Walker 2006), is also predominantly neutral and negative (Fig. S3). Additionally, only 3 of the 12,070 genes in this analysis showed statistically significant (p < 0.05) signatures of positive selection after multiple test corrections: CDHR2 (*Cadherin-related family member 2*, p = 0.00898), ITGB3 (*Integrin beta-3*, p = 0.00564), and HELZ2 (*Helicase with zinc finger domain 2*, p = 0.0404).

**Figure 3.**
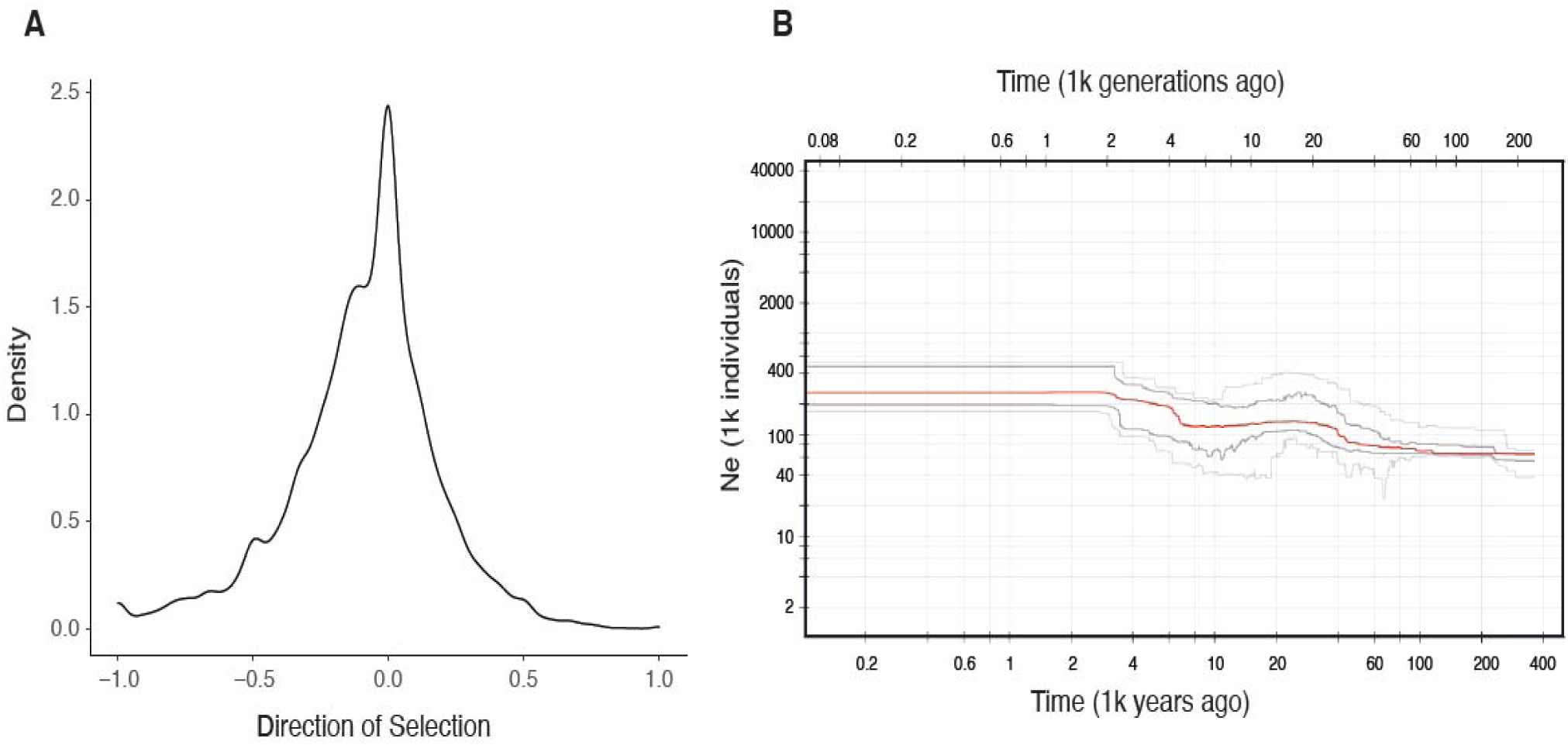
Molecular evolution of the black-headed duck. A) Distribution of the estimates of the direction of selection in coding regions and B) the inferred demography showing a relatively stable effective population size.

In both the GO enrichment analysis and the GSE analysis using the *G. gallus* database, there were no significantly enriched terms associated with *H. atricapilla*. However in the GSE analysis with the *Homo sapiens* database, there were three gene sets significantly associated with chemokine receptor activity (p_FDR_ < 0.1), which are required for the development and homeostasis of the immune system (Hughes & Nibbs, 2018).

Inferring the demography of *H. atricapilla* from the site frequency spectrum suggests a small population expansion occurred between 4,000 and 7,000 years ago, with the population size remaining stable until present day with an effective population size of approximately 250,000 individuals (Fig. 3B). Additionally, there is no evidence for selective sweeps on the major chromosomes of *H. atricapilla*.

## Discussion

Using newly assembled and annotated genomes, we investigated the genomic signals that are unique to the obligate brood parasite, *Heteronetta atricapilla*, in both coding and noncoding sequences in comparison to three closely related nesting species of waterfowl, and resolved the phylogenetic polytomy that existed between these four focal taxa (Fig. 1). Our *a priori* hypothesis was that there would be signatures of selection in protein-coding regions, non-coding regulatory regions, and/or selective sweeps across the genome. We identified 294 accelerated conserved non-exonic elements (CNEEs) that are unique to the *H. atricapilla* lineage (Table SX) which were associated with signals of neuronal development and olfactory receptor activity (Fig. 2, Table S3). We also identified three genes with evidence for significant positive selection, but overall observed predominantly negative and neutral selection in coding regions. While the presence of segregating deleterious mutations could be caused by a small or declining population size, we did not see any evidence to support this in our inference of demographic history (Fig. 3), nor did we observe any strong evidence for recent selective sweeps.

The number of accelerated CNEEs uniquely identified in each focal taxa was similar across species, but both *H. atricapilla* and *Nettapus auritus* had approximately 100 more accelerated elements than *Oxyura jamaicensis* and *Stictonetta naevosa* (Table SX). *N. auritus* is more distantly related to the other three focal species, diverging from the three ducks as a sister species in the coding, non-coding, and mitogenome tree reconstructions (Fig. 1). Within the three focal duck species, there were more accelerated elements that are uniquely accelerated in *H. atricapilla* than in the nesting species *O. jamaicensis* and *S. naevosa* (Table SX). However, the number of accelerated elements in each focal species is substantially less than other studies that have identified lineage-specific rate accelerations in CNEEs using avian species (e.g., Sackton et al. 2019). Additionally, the species we have sampled here are closely related, and thus it is not surprising that we did not identify a large set of uniquely accelerated elements for each species.

From the small set of CNEEs that were uniquely identified in *H. atricapilla*, there was a marginally significant peak of accelerated CNEEs on the Z chromosome, but no large clusters that would indicate spatial enrichment of accelerated elements (Fig. 2A). However, the predominant signals arising from the functional annotation of significantly enriched regions flanking genes, and associated GO terms, are those of neuronal development and olfactory receptor activity (Fig. 2, Table 1). The strong signals of accelerated evolution in putative regulatory regions of genes associated with neuronal/nervous system development could be attributed to the extreme precociality of the *H. atricapilla* chicks (Davies 2010). The chicks of *H. atricapilla* are even more independent and precocial than the typical nesting waterfowl, which is a notable divergence from the other avian obligate brood parasites and their altricial chicks (Lyon & Eadie 1991).

The signals of olfactory receptor activity observed in the GO term analysis are associated with OLFR6, an olfactory receptor. There has been a widely held belief that olfaction has not been an important sense in birds, or at the very least it is underdeveloped (Balthazart & Taziaux 2009). The foundational work of Michelsen (Michelsen 1959) on odour recognition in pigeons led to more of a focus on the role of olfaction in birds, with the work of Bang (Bang 1960) and Bang and Cobb (Bang & Cobb 1968) on the anatomy and variation of olfactory cavities and bulb sizes. There has been increasing recognition that olfaction can play an important role in many avian species (reviewed in Balthazart & Taziaux 2009) and genomic investigations have observed a highly localised region of the genome that contains hundreds of olfactory receptors in *G. gallus* (Driver & Balakrishnan 2021). Birds do possess a fully developed nasal cavity and recent work has highlighted the importance of the sensory perception of smell in a variety of tasks from food location (Castro et al. 2010) to mate choice (Balthazart & Taziaux 2009; Caro et al. 2015).

Furthermore, there is a trend for some avian species (including ducks) to have larger olfactory bulbs (Bang & Cobb 1968; Wenzel 1971) and olfaction plays a large role in male mallard reproductive success (Balthazart & Schoffeniels 1979; Caro & Balthazart 2010). While this olfactory signal could be associated with navigation in a semi-aquatic environment (Corfield et al. 2015), we only observe this signal in the brood parasitic *H. atricapilla* and not in the closely related nesting species of waterfowl. Given the breeding grounds of *H. atricapilla* are typically freshwater marsh environments (Cabrera et al. 2017) and that olfaction for nest localisation is prevalent in seabirds (Bonadonna et al. 2003; Bonadonna & Nevitt 2004), it could be hypothesised that olfaction may be important for this obligate brood parasite to find host nests in a densely vegetated habitat, but further work would be needed to support this.

In protein-coding regions, the distribution of signatures of selection on a per-gene basis was predominantly neutral and negative in both the direction of selection (Fig. 3A) and alpha (Fig. S3) calculations. We also only identified 19 genes that had statistically significant signatures of selection, and only three of those genes were positively selected for: CDHR2 (*Cadherin-related family member 2*), ITGB3 (*Integrin beta-3*), and HELZ2 (*Helicase with zinc finger domain 2*). CDHR2, which is also enriched for accelerated CNEEs in regulatory regions of *N. auritus*, creates calcium-dependent adhesion complexes with CDHR5 (*Cadherin-related family member 5*) on adjacent microvilli, controlling the packing of microvilli on epithelial cells along the brush border in humans (Crawley et al. 2014). Identifying signatures of selection in ITGB3 is not surprising, as immune-related genes are frequently under selection in many species (Shultz & Sackton 2019). HELZ2 is a transcriptional coactivator that works with THRAP3 (*Thyroid hormone receptor-associated protein 3*; Katano-Toki et al. 2013) to enhance the transcriptional activation of PPAR-Y, a master regulator of adipose tissue differentiation (Ahmadian et al. 2013). While these signals of positive selection could be worthy of further investigation in experimental manipulations, they are not associated with the phenotypes of obligate brood parasitism, notably in this lineage, the loss of parental care, nest building behaviours, chick provisioning, etc.

The evidence for segregating deleterious mutations in protein-coding regions could be explained by a small or declining population size in *H. atricapilla*, but we did not find any evidence to support this hypothesis from our inferred demographic history. We instead see a relatively constant effective population size of approximately 250,000 individuals following a small population expansion between 4,000 and 8,000 years ago (Fig. 3B), likely as a result of a post-glacial population expansion. While an effective population size of 250,000 individuals is not necessarily large, it is also not a small or declining population, and therefore is likely not the driving factor behind the mostly neutral and negative selection we observe in coding regions of *H. atricapilla*.

Overall, we do not observe clear signals of selection in *Heteronetta atricapilla* that are specifically associated with our *a priori* hypothesis concerning the loss of parental care. This is possibly because *H. atricapilla* is more of a generalist obligate brood parasite than the other avian lineages. Additionally, *H. atricapilla* is likely constrained from evolving in many brood parasitic traits due to the aggressive intraspecific brood parasitism that occurs in its main host species (Lyon & Eadie 1991). Ultimately, intraspecific and facultative brood parasitism is relatively common in waterfowl, which may also contribute to a lack of specific signals in *H. atricapilla* related to obligate brood parasitism. However, we have identified more signals of selection in non-coding regulatory regions than in coding sequences, of which these selective pressures may be associated with development of extremely precocial chicks or olfactory receptors that could be important in nest searching.

## Acknowledgements

The authors would like to thank Dustin Foote and the Sylvan Heights Bird Park for providing *Heteronetta atricapilla* samples for the population resequencing data. The computations in this paper were run on the FASRC Odyssey cluster supported by the FAS Division of Science Research Computing Group at Harvard University. This work was funded by a National Science Foundation grant awarded to MDS to research the comparative genomics of host-specific adaptation and life history evolution in brood parasitic birds.

## Notes

### Competing Interest Statement

The authors have declared no competing interest.

## Literature Cited

Ahmadian M et al. 2013. PPARγ signaling and metabolism: the good, the bad and the future. Nat. Med. 19:557–566.

Armstrong J et al. 2020. Progressive Cactus is a multiple-genome aligner for the thousand-genome era. Nature. 587:246–251.

Balthazart J, Schoffeniels E. 1979. Pheromones are involved in the control of sexual behaviour in birds. Naturwissenschaften. 66:55–56.

Balthazart J, Taziaux M. 2009. The underestimated role of olfaction in avian reproduction? Behav. Brain Res. 200:248–259.

Bang BG. 1960. Anatomical evidence for olfactory function in some species of birds. Nature. 188:547–549.

Bang BG, Cobb S. 1968. The Size of the Olfactory Bulb in 108 Species of Birds. Auk. 85:55–61.

Bonadonna F, Cunningham GB, Jouventin P, Hesters F, Nevitt GA. 2003. Evidence for nest-odour recognition in two species of diving petrel. J. Exp. Biol. 206:3719–3722.

Bonadonna F, Nevitt GA. 2004. Partner-specific odor recognition in an Antarctic seabird. Science. 306:835.

Bouckaert R et al. 2019. BEAST 2.5: An advanced software platform for Bayesian evolutionary analysis. PLoS Comput. Biol. 15:e1006650.

Cabrera M, Montalti D, Segura L. 2017. Breeding Phenology and New Host List of the Black-Headed Duck (Heteronetta atricapilla) In Argentina. The Wilson Journal of Ornithology. 129:311–316.

Caro SP, Balthazart J. 2010. Pheromones in birds: myth or reality? Journal of Comparative Physiology A. 196:751–766.

Caro SP, Balthazart J, Bonadonna F. 2015. The perfume of reproduction in birds: Chemosignaling in avian social life. Horm. Behav. 68:25–42.

Castro I et al. 2010. Olfaction in birds: a closer look at the kiwi (Apterygidae). J. Avian Biol. 41:213–218.

Cingolani P et al. 2012. A program for annotating and predicting the effects of single nucleotide polymorphisms, SnpEff: SNPs in the genome of Drosophila melanogaster strain w1118; iso-2; iso-3. Fly. 6:80–92.

Corfield JR et al. 2015. Diversity in olfactory bulb size in birds reflects allometry, ecology, and phylogeny. Front. Neuroanat. 9:102.

Craig RJ, Suh A, Wang M, Ellegren H. 2018. Natural selection beyond genes: Identification and analyses of evolutionarily conserved elements in the genome of the collared flycatcher (Ficedula albicollis). Mol. Ecol. 27:476–492.

Crawley SW et al. 2014. Intestinal brush border assembly driven by protocadherin-based intermicrovillar adhesion. Cell. 157:433–446.

Davies N. 2010. Cuckoos, Cowbirds and Other Cheats. A&C Black.

DeGiorgio M, Huber CD, Hubisz MJ, Hellmann I, Nielsen R. 2016. SweepFinder2: increased sensitivity, robustness and flexibility. Bioinformatics. 32:1895–1897.

Di Franco A, Poujol R, Baurain D, Philippe H. 2019. Evaluating the usefulness of alignment filtering methods to reduce the impact of errors on evolutionary inferences. BMC Evol. Biol. 19:1–17.

Driver RJ, Balakrishnan CN. 2021. Highly Contiguous Genomes Improve the Understanding of Avian Olfactory Receptor Repertoires. Integr. Comp. Biol. doi: 10.1093/icb/icab150.

Durinck S, Spellman PT, Birney E, Huber W. 2009. Mapping identifiers for the integration of genomic datasets with the R/Bioconductor package biomaRt. Nat. Protoc. 4:1184–1191.

Edgar RC. 2004. MUSCLE: a multiple sequence alignment method with reduced time and space complexity. BMC Bioinformatics. 5:1–19.

Eilertson KE, Booth JG, Bustamante CD. 2012. SnIPRE: Selection Inference Using a Poisson Random Effects Model. PLoS Computational Biology. 8:e1002806. doi: 10.1371/journal.pcbi.1002806.

Emms DM, Kelly S. 2019. OrthoFinder: phylogenetic orthology inference for comparative genomics. Genome Biol. 20:238.

Eyre-Walker A. 2006. The genomic rate of adaptive evolution. Trends Ecol. Evol. 21:569–575.

Feeney WE, Welbergen JA, Langmore NE. 2012. The frontline of avian brood parasite-host coevolution. Anim. Behav. 84:3–12.

Gonzalez J, Düttmann H, Wink M. 2009. Phylogenetic relationships based on two mitochondrial genes and hybridization patterns in Anatidae. J. Zool.. 279:310–318.

Hackett SJ et al. 2008. A Phylogenomic Study of Birds Reveals Their Evolutionary History. Science. 320:1763–1768.

Hickey G, Paten B, Earl D, Zerbino D, Haussler D. 2013. HAL: a hierarchical format for storing and analyzing multiple genome alignments. Bioinformatics. 29:1341–1342.

Hubisz MJ, Pollard KS, Siepel A. 2011. PHAST and RPHAST: phylogenetic analysis with space/time models. Brief. Bioinform. 12:41–51.

Hughes CE, and Nibbs RJB. 2018. A guide to chemokines and their receptors. FEBS J. 285: 2944–2971.

Hu Z, Sackton TB, Edwards SV, Liu JS. 2019. Bayesian Detection of Convergent Rate Changes of Conserved Noncoding Elements on Phylogenetic Trees. Mol. Biol. Evol. 36:1086–1100.

Junier T, Zdobnov EM. 2010. The Newick utilities: high-throughput phylogenetic tree processing in the UNIX shell. Bioinformatics. 26:1669–1670.

Katano-Toki A et al. 2013. THRAP3 Interacts with HELZ2 and Plays a Novel Role in Adipocyte Differentiation. Mol. Endocrinol. 27:769–780.

Katoh K, Standley DM. 2013. MAFFT multiple sequence alignment software version 7: improvements in performance and usability. Mol. Biol. Evol. 30:772–780.

Kearse M et al. 2012. Geneious Basic: an integrated and extendable desktop software platform for the organization and analysis of sequence data. Bioinformatics. 28:1647–1649.

König S, Romoth LW, Gerischer L, Stanke M. 2016. Simultaneous gene finding in multiple genomes. Bioinformatics. 32:3388–3395.

Kosakovsky Pond SL, Frost SDW. 2005. Not so different after all: a comparison of methods for detecting amino acid sites under selection. Mol. Biol. Evol. 22:1208–1222.

Kozlov AM, Darriba D, Flouri T, Morel B, Stamatakis A. 2019. RAxML-NG: a fast, scalable and user-friendly tool for maximum likelihood phylogenetic inference. Bioinformatics. 35:4453–4455.

Lanfear R, Frandsen PB, Wright AM, Senfeld T, Calcott B. 2016. PartitionFinder 2: New Methods for Selecting Partitioned Models of Evolution for Molecular and Morphological Phylogenetic Analyses. Mol. Biol. Evol. 34:772–773.

Liu X, Fu Y-X. 2020. Author Correction: Stairway Plot 2: demographic history inference with folded SNP frequency spectra. Genome Biol. 21:305.

Livezey B. 1995. Phylogeny and comparative ecology of stiff-tailed ducks (Anatidae: Oxyurini). https://www.semanticscholar.org/paper/850170e4f1d7094938a0dcdbd792fe640c9fbe33 (Accessed August 16, 2021).

Lowe CB, Clarke JA, Baker AJ, Haussler D, Edwards SV. 2014. Feather Development Genes and Associated Regulatory Innovation Predate the Origin of Dinosauria. Mol. Biol. Evol. 32:23–28.

Lynch KS, O’Connell LA, Louder MIM, Balakrishnan CN, Fischer EK. 2019. Understanding the Loss of Maternal Care in Avian Brood Parasites Using Preoptic Area Transcriptome Comparisons in Brood Parasitic and Non-parasitic Blackbirds. G3: Genes, Genomes, Genetics. 9:1075–1084.

Lyon BE, Eadie JM. 2004. An obligate brood parasite trapped in the intraspecific arms race of its hosts. Nature. 432:390–393.

Lyon BE, Eadie JM. 1991. Mode of development and interspecific avian brood parasitism. Behav. Ecol. 2:309–318.

Lyon BE, Eadie JM. 2013. Patterns of host use by a precocial obligate brood parasite, the Black-headed Duck: ecological and evolutionary considerations. Chin. Birds. 4:71–85.

McDonald JH, Kreitman M. 1991. Adaptive protein evolution at the Adh locus in Drosophila. Nature. 351:652–654.

Michelsen WJ. 1959. Procedure for Studying Olfactory Discrimination in Pigeons. Science. 130:630–631.

Murrell B et al. 2015. Gene-wide identification of episodic selection. Mol. Biol. Evol. 32:1365–1371.

Payne RB. 1974. The Evolution of Clutch Size and Reproductive Rates in Parasitic Cuckoos. Evolution. 28:169–181.

Pertea G, Pertea M. 2020. GFF Utilities: GffRead and GffCompare. F1000 Res. 9: 304.

Pond SLK, Frost SDW, Muse SV. 2004. HyPhy: hypothesis testing using phylogenies. Bioinformatics. 21:676–679.

Rees E, Hillgarth N. 1984. The breeding biology of captive black-headed ducks and the behavior of their yound. Condor. 242–250.

Roldán M, Soler M. 2011. Parental-care parasitism: how do unrelated offspring attain acceptance by foster parents? Behav. Ecol. 22:679–691.

Rothstein SI. 1990. A Model System for Coevolution: Avian Brood Parasitism. Annu. Rev. Ecol. Syst. 21:481–508.

Sackton TB et al. 2019. Convergent regulatory evolution and loss of flight in paleognathous birds. Science. 364:74–78.

Schneider ER et al. 2019. A Cross-Species Analysis Reveals a General Role for Piezo2 in Mechanosensory Specialization of Trigeminal Ganglia from Tactile Specialist Birds. Cell Rep. 26:1979–1987.e3.

Scott DM, Ankney CD. 1983. The laying cycle of Brown-headed Cowbirds: Passerine chickens? Auk. 100:583–592.

Seppey M, Manni M, Zdobnov EM. 2019. BUSCO: Assessing Genome Assembly and Annotation Completeness. Methods Mol. Biol. 1962:227–245.

Sherry DF, Guigueno MF. 2019. Cognition and the brain of brood parasitic cowbirds. Integr. Zool. 14:145–157.

Shultz AJ, Sackton TB. 2019. Immune genes are hotspots of shared positive selection across birds and mammals. Elife. 8:e41815.

Siepel A et al. 2005. Evolutionarily conserved elements in vertebrate, insect, worm, and yeast genomes. Genome Res. 15:1034–1050.

Smeds L, Qvarnström A, Ellegren H. 2016. Direct estimate of the rate of germline mutation in a bird. Genome Res. 26:1211–1218.

Smith MD et al. 2015. Less is more: an adaptive branch-site random effects model for efficient detection of episodic diversifying selection. Mol. Biol. Evol. 32:1342–1353.

Soler M, ed. 2017. Avian Brood Parasitism: Behaviour, Ecology, Evolution and Coevolution. Springer, Cham.

Sorenson MD, Payne RB. 2002. Molecular Genetic Perspectives on Avian Brood Parasitism1. Integr. Comp. Biol. 42:388–400.

Stanke M, Diekhans M, Baertsch R, Haussler D. 2008. Using native and syntenically mapped cDNA alignments to improve de novo gene finding. Bioinformatics. 24:637–644.

Stoddard MC, Hauber ME. 2017. Colour, vision and coevolution in avian brood parasitism. Philos. Trans. R. Soc. Lond. B Biol. Sci. 372. doi: 10.1098/rstb.2016.0339.

Stoletzki N, Eyre-Walker A. 2011. Estimation of the Neutrality Index. Molecular Biology and Evolution. 28:63–70. doi: 10.1093/molbev/msq249.

Sun Z et al. 2017. Rapid and recent diversification patterns in Anseriformes birds: Inferred from molecular phylogeny and diversification analyses. PLoS One. 12:e0184529.

Van der Auwera GA, O’Connor BD. 2020. Genomics in the Cloud: Using Docker, GATK, and WDL in Terra. ‘O’Reilly Media, Inc.’

Weisenfeld NI, Kumar V, Shah P, Church DM, Jaffe DB. 2017. Direct determination of diploid genome sequences. Genome Res. 27:757–767.

Weller MW. 1968. The breeding biology of the black-headed duck. Living Bird. 7:169–208.

Wenzel BM. 1971. Olfactory sensation in the kiwi and other birds. Ann. N. Y. Acad. Sci. 188:183–193.

Yu G, Wang L-G, Han Y, He Q-Y. 2012. clusterProfiler: an R package for comparing biological themes among gene clusters. OMICS. 16:284–287.

Zhang C, Rabiee M, Sayyari E, Mirarab S. 2018. ASTRAL-III: polynomial time species tree reconstruction from partially resolved gene trees. BMC Bioinformatics. 19:15–30.

