## Supplemental Figure 1 for "Patterns of lineage-specific genome evolution in the brood parasitic black-headed duck (*Heteronetta atricapilla*)"

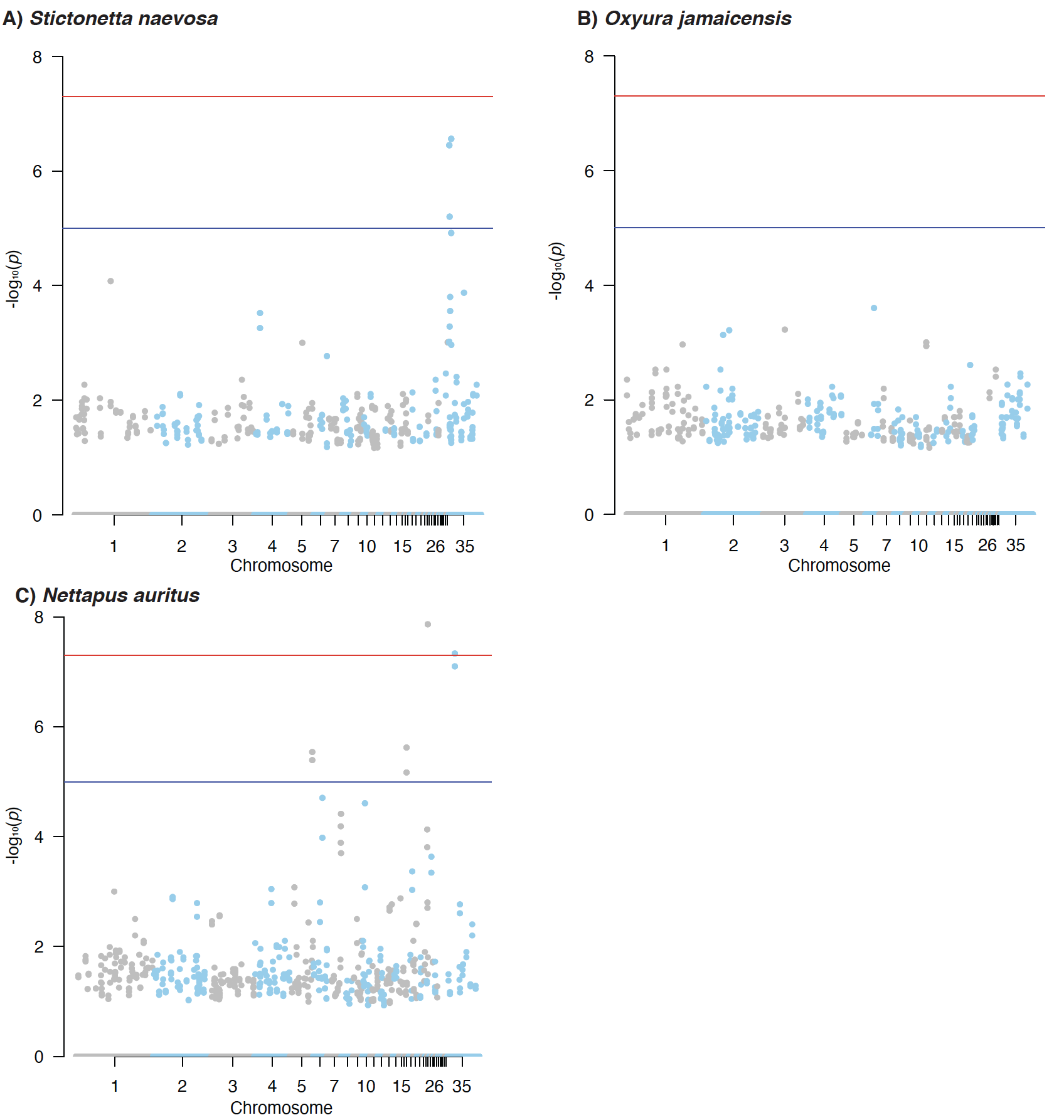

**Figure S1.** Genome-wide scans for clusters of accelerated conserved non-exonic elements (CNEEs) in A) freckled duck, B) ruddy duck, and C) African pygmy goose.
