## Supplemental Figure 2 for "Patterns of lineage-specific genome evolution in the brood parasitic black-headed duck (*Heteronetta atricapilla*)"

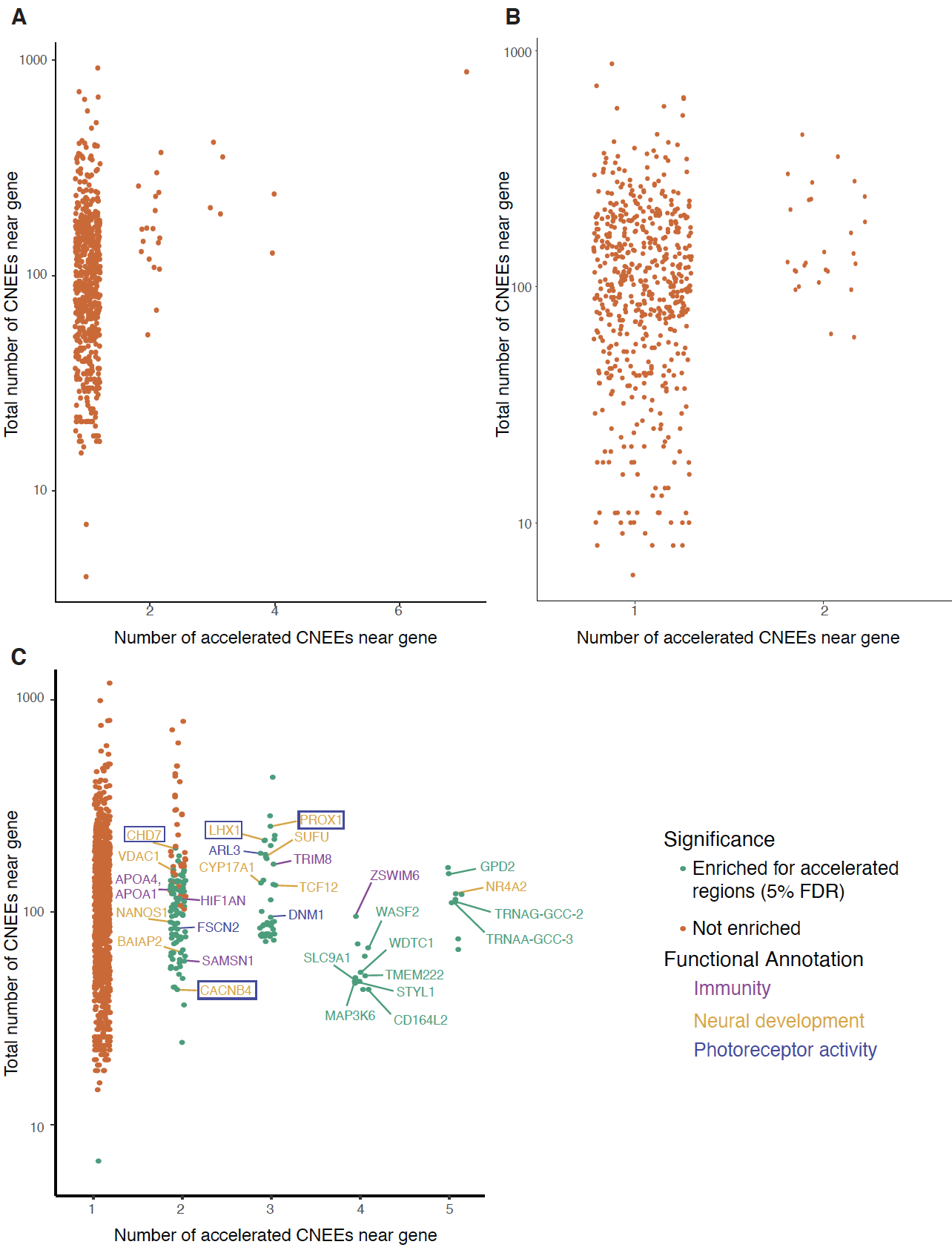

**Figure S2**. Genes significantly enriched for accelerated regions of CNEEs in A) freckled duck, B) ruddy duck, and C) African pygmy goose.
