## Supplemental Figure 3 for "Patterns of lineage-specific genome evolution in the brood parasitic black-headed duck (*Heteronetta atricapilla*)"

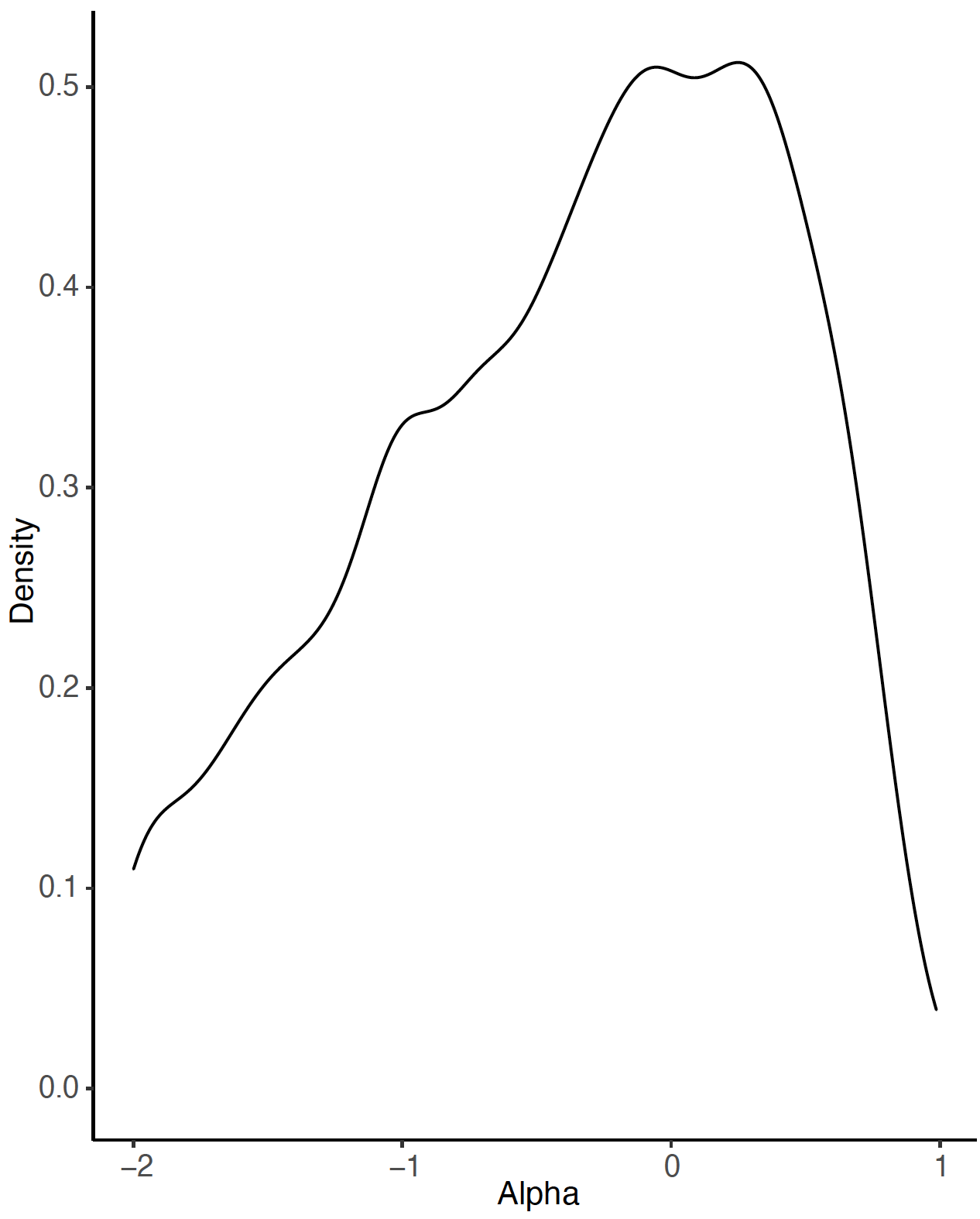

**Figure S3.** Distribution of alpha on a per gene basis in *Heteronetta atricapilla* (the proportion of amino acids fixed by positive selection).
