## Supplemental Table 1 for "Patterns of lineage-specific genome evolution in the brood parasitic black-headed duck (*Heteronetta atricapilla*)"

**Table S1.** Descriptive statistics of genome assemblies and annotations of focal Anseriformes species.

| Species | Raw Coverage | Contig N50 (Kb) | Scaffold N50 (Mb) | Assembly Size (Gb) | Assembly in Scaffolds >10Kb (%) | Genome Assembly Completeness (%,  BUSCO C score) | Genome Annotation Completeness (%,  BUSCO C score) |
| --- | --- | --- | --- | --- | --- | --- | --- |
| *Heteronetta atricapilla* | 53.17 | 155.79 | 23.17 | 1.1 | 96.74 | 94.7 | 91 |
| *Stictonetta naevosa* | 41.7 | 142.23 | 10.96 | 1.06 | 96.5 | 95 | 90.6 |
| *Oxyura jamaicensis* | 54.96 | 132.48 | 46.21 | 1.09 | 96.15 | 94.7 | 92.6 |
| *Nettapus auritus* | 50.2 | 108.05 | 25.86 | 1.09 | 95.84 | 94.6 | 91.3 |
