## Supplemental Table 2 for "Patterns of lineage-specific genome evolution in the brood parasitic black-headed duck (*Heteronetta atricapilla*)"

**Table S2.** Publicly available genome assemblies used in the whole genome alignment and downstream comparative analyses.

| **Species** | **Common Name** | **GenBank Accession** |
| --- | --- | --- |
| Anas platyrhynchos | Mallard | GCA_003850225.1 |
| Colinus virginianus | Northern bobwhite | GCA_000599465.2 |
| Gallus gallus | Chicken (red jungle fowl) | GCA_000002315.5 |
| Branta canadensis | Canada goose | GCA_006130075.1 |
| Tympanuchus cupido pinnatus | Greater prairie chicken | GCA_001870855.1 |
| Syrmaticus mikado | Mikado pheasant | GCA_003435085.1 |
| Anser indicus | Bar-headed goose | GCA_006229135.1 |
| Numida meleagris | Helmeted guineafowl | GCA_002078875.2 |
| Anser brachyrhynchus | Pink-footed goose | GCA_002592135.1 |
| Coturnix japonica | Japanese quail | GCA_001577835.2 |
| Anser cygnoides domesticus | Swan goose | GCA_002166845.1 |
| Heteronetta atricapilla | Black-headed duck | GCA_011075105.1 |
| Nettapus auritus | African pygmy goose | GCA_011076525.1 |
| Oxyura jamaicensis | Ruddy duck | GCA_011077185.1 |
| Stictonetta naevosa | Freckled duck | GCA_011074415.1 |
