## Supplemental Table 3 for "Patterns of lineage-specific genome evolution in the brood parasitic black-headed duck (*Heteronetta atricapilla*)"

**Table S3.** Gene Ontology (GO) terms in nesting focal species.

| Species | GO Term | p-value | Subontology | Functional Annotation |
| --- | --- | --- | --- | --- |
| *Stictonetta naevosa* | GO:0022607 | 0.001998 | BP | cellular component assembly |
| *Stictonetta naevosa* | GO:0044085 | 0.00499501 | BP | cellular component biogenesis |
| *Stictonetta naevosa* | GO:0043933 | 0.01198801 | BP | protein-containing complex subunit organization |
| *Stictonetta naevosa* | GO:0018193 | 0.01698302 | BP | peptidyl-amino acid modification |
| *Stictonetta naevosa* | GO:0071495 | 0.01798202 | BP | cellular response to endogenous stimulus |
| *Stictonetta naevosa* | GO:0007417 | 0.01998002 | BP | central nervous system development |
| *Stictonetta naevosa* | GO:0048468 | 0.01998002 | BP | cell development |
| *Stictonetta naevosa* | GO:0048869 | 0.02997003 | BP | cellular developmental process |
| *Stictonetta naevosa* | GO:0051130 | 0.03196803 | BP | positive regulation of cellular component organization |
| *Stictonetta naevosa* | GO:0030154 | 0.03596404 | BP | cell differentiation |
| *Stictonetta naevosa* | GO:0070848 | 0.03796204 | BP | response to growth factor |
| *Stictonetta naevosa* | GO:0060429 | 0.03896104 | BP | epithelium development |
| *Stictonetta naevosa* | GO:0030054 | 0.01698302 | CC | cell junction |
| *Stictonetta naevosa* | GO:0012505 | 0.01998002 | CC | endomembrane system |
| *Stictonetta naevosa* | GO:0016020 | 0.02497502 | CC | membrane |
| *Stictonetta naevosa* | GO:0032991 | 0.03196803 | CC | protein-containing complex |
| *Stictonetta naevosa* | GO:0031224 | 0.04095904 | CC | intrinsic component of membrane |
| *Stictonetta naevosa* | GO:0038023 | 0.02897103 | MF | signaling receptor activity |
| *Stictonetta naevosa* | GO:0060089 | 0.02897103 | MF | molecular transducer activity |
| *Oxyura jamaicensis* | GO:0007010 | 0.00999001 | BP | cytoskeleton organization |
| *Oxyura jamaicensis* | GO:0071310 | 0.01298701 | BP | cellular response to organic substance |
| *Oxyura jamaicensis* | GO:0071495 | 0.02297702 | BP | cellular response to endogenous stimulus |
| *Oxyura jamaicensis* | GO:0044093 | 0.02597403 | BP | positive regulation of molecular function |
| *Oxyura jamaicensis* | GO:0051336 | 0.02597403 | BP | regulation of hydrolase activity |
| *Oxyura jamaicensis* | GO:0006996 | 0.02697303 | BP | organelle organization |
| *Oxyura jamaicensis* | GO:0033043 | 0.03396603 | BP | regulation of organelle organization |
| *Oxyura jamaicensis* | GO:0009719 | 0.03496504 | BP | response to endogenous stimulus |
| *Oxyura jamaicensis* | GO:0030029 | 0.03496504 | BP | actin filament-based process |
| *Oxyura jamaicensis* | GO:0071363 | 0.03496504 | BP | cellular response to growth factor stimulus |
| *Oxyura jamaicensis* | GO:0030036 | 0.04395604 | BP | actin cytoskeleton organization |
| *Oxyura jamaicensis* | GO:0034622 | 0.04495505 | BP | cellular protein-containing complex assembly |
| *Oxyura jamaicensis* | GO:0044087 | 0.04895105 | BP | regulation of cellular component biogenesis |
| *Oxyura jamaicensis* | GO:1902494 | 0.03496504 | CC | catalytic complex |
| *Oxyura jamaicensis* | GO:0030054 | 0.03696304 | CC | cell junction |
| *Oxyura jamaicensis* | GO:0005833 | 0.04195804 | CC | hemoglobin complex |
| *Oxyura jamaicensis* | GO:0031838 | 0.04195804 | CC | haptoglobin-hemoglobin complex |
| *Oxyura jamaicensis* | GO:0097367 | 0.02297702 | MF | carbohydrate derivative binding |
| *Oxyura jamaicensis* | GO:0031720 | 0.03696304 | MF | haptoglobin binding |
| *Oxyura jamaicensis* | GO:0044877 | 0.03896104 | MF | protein-containing complex binding |
| *Oxyura jamaicensis* | GO:0043168 | 0.04295704 | MF | anion binding |
| *Nettapus auritus* | GO:0006508 | 0.003996 | BP | proteolysis |
| *Nettapus auritus* | GO:1901701 | 0.00599401 | BP | cellular response to oxygen molecular entity |
| *Nettapus auritus* | GO:0050808 | 0.01398601 | BP | synapse organization |
| *Nettapus auritus* | GO:0002755 | 0.01798202 | BP | MyD88-dependent toll-like receptor signaling pathway |
| *Nettapus auritus* | GO:0007017 | 0.01998002 | BP | microtubule-based process |
| *Nettapus auritus* | GO:0048588 | 0.02097902 | BP | developmental cell growth |
| *Nettapus auritus* | GO:0010721 | 0.02397602 | BP | negative regulation of cell development |
| *Nettapus auritus* | GO:0060560 | 0.02397602 | BP | developmental growth involved in morphogenesis |
| *Nettapus auritus* | GO:0005975 | 0.02597403 | BP | carbohydrate metabolic process |
| *Nettapus auritus* | GO:0009084 | 0.02597403 | BP | glutamine family amino acid formation/biosynthesis/anabolism/synthesis |
| *Nettapus auritus* | GO:0150076 | 0.02597403 | BP | neuroinflammatory response |
| *Nettapus auritus* | GO:0031329 | 0.02797203 | BP | regulation of cellular catabolism/breakdown/degradation |
| *Nettapus auritus* | GO:0044257 | 0.02797203 | BP | cellular protein catabolic process |
| *Nettapus auritus* | GO:0050768 | 0.02797203 | BP | negative regulation of neurogenesis |
| *Nettapus auritus* | GO:0000226 | 0.02897103 | BP | microtubule cytoskeleton organization |
| *Nettapus auritus* | GO:0051961 | 0.02897103 | BP | down-regulation/inhibition of nervous system development |
| *Nettapus auritus* | GO:0006525 | 0.02997003 | BP | arginine metabolic process |
| *Nettapus auritus* | GO:0051603 | 0.02997003 | BP | proteolysis involved in cellular protein catabolic process |
| *Nettapus auritus* | GO:0002062 | 0.03096903 | BP | chondrocyte differentiation |
| *Nettapus auritus* | GO:0009620 | 0.03096903 | BP | response to fungus |
| *Nettapus auritus* | GO:0032787 | 0.03096903 | BP | monocarboxylic acid metabolic process |
| *Nettapus auritus* | GO:0034248 | 0.03096903 | BP | regulation of cellular amide metabolic process |
| *Nettapus auritus* | GO:0050770 | 0.03096903 | BP | regulation of axonogenesis |
| *Nettapus auritus* | GO:0043123 | 0.03196803 | BP | positive regulation of I-kappaB kinase/NF-kappaB signaling |
| *Nettapus auritus* | GO:0071407 | 0.03196803 | BP | cellular response to organic cyclic compound |
| *Nettapus auritus* | GO:0072073 | 0.03196803 | BP | kidney epithelium development |
| *Nettapus auritus* | GO:0002224 | 0.03396603 | BP | toll-like receptor signaling pathway |
| *Nettapus auritus* | GO:0018205 | 0.03396603 | BP | peptidyl-lysine modification |
| *Nettapus auritus* | GO:0002886 | 0.03496504 | BP | regulation of myeloid leukocyte mediated immunity |
| *Nettapus auritus* | GO:0071103 | 0.03496504 | BP | DNA conformation change |
| *Nettapus auritus* | GO:0090287 | 0.03496504 | BP | regulation of cellular response to growth factor stimulus |
| *Nettapus auritus* | GO:0140115 | 0.03496504 | BP | export across plasma membrane |
| *Nettapus auritus* | GO:1905114 | 0.03496504 | BP | cell surface receptor signaling pathway involved in cell-cell signaling |
| *Nettapus auritus* | GO:0006418 | 0.03696304 | BP | tRNA aminoacylation for protein translation |
| *Nettapus auritus* | GO:0019405 | 0.03696304 | BP | alditol catabolic process |
| *Nettapus auritus* | GO:0022616 | 0.03696304 | BP | DNA strand elongation |
| *Nettapus auritus* | GO:0030111 | 0.03696304 | BP | regulation of Wnt signaling pathway |
| *Nettapus auritus* | GO:0043038 | 0.03696304 | BP | amino acid activation |
| *Nettapus auritus* | GO:0043039 | 0.03696304 | BP | tRNA aminoacylation |
| *Nettapus auritus* | GO:0044275 | 0.03696304 | BP | cellular carbohydrate catabolic process |
| *Nettapus auritus* | GO:0045926 | 0.03896104 | BP | negative regulation of growth |
| *Nettapus auritus* | GO:0070207 | 0.03996004 | BP | protein homotrimerization |
| *Nettapus auritus* | GO:0070371 | 0.03996004 | BP | ERK1 and ERK2 cascade |
| *Nettapus auritus* | GO:0001823 | 0.04095904 | BP | mesonephros development |
| *Nettapus auritus* | GO:0032479 | 0.04095904 | BP | regulation of type I interferon production |
| *Nettapus auritus* | GO:0072163 | 0.04095904 | BP | mesonephric epithelium development |
| *Nettapus auritus* | GO:0072164 | 0.04095904 | BP | mesonephric tubule development |
| *Nettapus auritus* | GO:0001959 | 0.04195804 | BP | regulation of cytokine-mediated signaling pathway |
| *Nettapus auritus* | GO:0034763 | 0.04195804 | BP | negative regulation of transmembrane transport |
| *Nettapus auritus* | GO:0043618 | 0.04195804 | BP | regulation of transcription from RNA polymerase II promoter in response to stress |
| *Nettapus auritus* | GO:0043620 | 0.04195804 | BP | regulation of DNA-templated transcription in response to stress |
| *Nettapus auritus* | GO:0060759 | 0.04195804 | BP | regulation of response to cytokine stimulus |
| *Nettapus auritus* | GO:0070897 | 0.04195804 | BP | transcription preinitiation complex assembly |
| *Nettapus auritus* | GO:0003009 | 0.04295704 | BP | skeletal muscle contraction |
| *Nettapus auritus* | GO:0019941 | 0.04295704 | BP | modification-dependent protein catabolic process |
| *Nettapus auritus* | GO:0038061 | 0.04295704 | BP | NIK/NF-kappaB signaling |
| *Nettapus auritus* | GO:0043632 | 0.04295704 | BP | modification-dependent macromolecule catabolic process |
| *Nettapus auritus* | GO:0048511 | 0.04295704 | BP | rhythmic process |
| *Nettapus auritus* | GO:0050879 | 0.04295704 | BP | multicellular organismal movement |
| *Nettapus auritus* | GO:0050881 | 0.04295704 | BP | musculoskeletal movement |
| *Nettapus auritus* | GO:2001243 | 0.04295704 | BP | negative regulation of intrinsic apoptotic signaling pathway |
| *Nettapus auritus* | GO:0006511 | 0.04395604 | BP | ubiquitin-dependent protein catabolic process |
| *Nettapus auritus* | GO:0022407 | 0.04395604 | BP | regulation of cell-cell adhesion |
| *Nettapus auritus* | GO:0045665 | 0.04395604 | BP | negative regulation of neuron differentiation |
| *Nettapus auritus* | GO:0060828 | 0.04395604 | BP | regulation of canonical Wnt signaling pathway |
| *Nettapus auritus* | GO:0070206 | 0.04395604 | BP | protein trimerization |
| *Nettapus auritus* | GO:1990138 | 0.04395604 | BP | neuron projection extension |
| *Nettapus auritus* | GO:0000387 | 0.04495505 | BP | spliceosomal snRNP assembly |
| *Nettapus auritus* | GO:0051865 | 0.04495505 | BP | protein autoubiquitination |
| *Nettapus auritus* | GO:0008361 | 0.04595405 | BP | regulation of cell size |
| *Nettapus auritus* | GO:0009584 | 0.04595405 | BP | detection of visible light |
| *Nettapus auritus* | GO:1901652 | 0.04595405 | BP | response to peptide |
| *Nettapus auritus* | GO:0001732 | 0.04695305 | BP | formation of cytoplasmic translation initiation complex |
| *Nettapus auritus* | GO:0006368 | 0.04695305 | BP | transcription elongation from RNA polymerase II promoter |
| *Nettapus auritus* | GO:0031440 | 0.04695305 | BP | regulation of mRNA 3'-end processing |
| *Nettapus auritus* | GO:0042558 | 0.04695305 | BP | pteridine-containing compound metabolic process |
| *Nettapus auritus* | GO:0046164 | 0.04695305 | BP | alcohol catabolic process |
| *Nettapus auritus* | GO:0046174 | 0.04695305 | BP | polyol catabolic process |
| *Nettapus auritus* | GO:0061387 | 0.04695305 | BP | regulation of extent of cell growth |
| *Nettapus auritus* | GO:0097503 | 0.04695305 | BP | sialylation |
| *Nettapus auritus* | GO:0006417 | 0.04795205 | BP | regulation of translation |
| *Nettapus auritus* | GO:0006914 | 0.04795205 | BP | autophagy |
| *Nettapus auritus* | GO:0007602 | 0.04795205 | BP | phototransduction |
| *Nettapus auritus* | GO:0016055 | 0.04795205 | BP | Wnt signaling pathway |
| *Nettapus auritus* | GO:0031397 | 0.04795205 | BP | negative regulation of protein ubiquitination |
| *Nettapus auritus* | GO:0036473 | 0.04795205 | BP | cell death in response to oxidative stress |
| *Nettapus auritus* | GO:0042060 | 0.04795205 | BP | wound healing |
| *Nettapus auritus* | GO:0061919 | 0.04795205 | BP | process utilizing autophagic mechanism |
| *Nettapus auritus* | GO:0198738 | 0.04795205 | BP | cell-cell signaling by wnt |
| *Nettapus auritus* | GO:1900180 | 0.04795205 | BP | regulation of protein localization to nucleus |
| *Nettapus auritus* | GO:1901881 | 0.04795205 | BP | positive regulation of protein depolymerization |
| *Nettapus auritus* | GO:1903321 | 0.04795205 | BP | negative regulation of protein modification by small protein conjugation or removal |
| *Nettapus auritus* | GO:0002279 | 0.04895105 | BP | mast cell activation involved in immune response |
| *Nettapus auritus* | GO:0002448 | 0.04895105 | BP | mast cell mediated immunity |
| *Nettapus auritus* | GO:0043299 | 0.04895105 | BP | leukocyte degranulation |
| *Nettapus auritus* | GO:0043303 | 0.04895105 | BP | mast cell degranulation |
| *Nettapus auritus* | GO:0045576 | 0.04895105 | BP | mast cell activation |
| *Nettapus auritus* | GO:0051259 | 0.04895105 | BP | protein complex oligomerization |
| *Nettapus auritus* | GO:0002088 | 0.04995005 | BP | lens development in camera-type eye |
| *Nettapus auritus* | GO:0009583 | 0.04995005 | BP | detection of light stimulus |
| *Nettapus auritus* | GO:0050892 | 0.04995005 | BP | intestinal absorption |
| *Nettapus auritus* | GO:0070372 | 0.04995005 | BP | regulation of ERK1 and ERK2 cascade |
| *Nettapus auritus* | GO:0097060 | 0.01198801 | CC | synaptic membrane |
| *Nettapus auritus* | GO:0000139 | 0.01998002 | CC | Golgi membrane |
| *Nettapus auritus* | GO:0005815 | 0.02897103 | CC | microtubule organizing center |
| *Nettapus auritus* | GO:0005911 | 0.03296703 | CC | cell-cell junction |
| *Nettapus auritus* | GO:0098590 | 0.03796204 | CC | plasma membrane region |
| *Nettapus auritus* | GO:0150034 | 0.04095904 | CC | distal axon |
| *Nettapus auritus* | GO:0005794 | 0.04195804 | CC | Golgi apparatus |
| *Nettapus auritus* | GO:0042555 | 0.04295704 | CC | MCM complex |
| *Nettapus auritus* | GO:0030532 | 0.04595405 | CC | small nuclear ribonucleoprotein complex |
| *Nettapus auritus* | GO:0044297 | 0.04595405 | CC | cell body |
| *Nettapus auritus* | GO:0097525 | 0.04595405 | CC | spliceosomal snRNP complex |
| *Nettapus auritus* | GO:0000815 | 0.04795205 | CC | ESCRT III complex |
| *Nettapus auritus* | GO:0016020 | 0.04795205 | CC | membrane |
| *Nettapus auritus* | GO:0036452 | 0.04795205 | CC | ESCRT complex |
| *Nettapus auritus* | GO:0005861 | 0.04995005 | CC | troponin complex |
| *Nettapus auritus* | GO:0043169 | 0.01698302 | MF | cation binding |
| *Nettapus auritus* | GO:0046872 | 0.01798202 | MF | metal ion binding |
| *Nettapus auritus* | GO:0005104 | 0.02297702 | MF | fibroblast growth factor receptor binding |
| *Nettapus auritus* | GO:0004620 | 0.02597403 | MF | phospholipase activity |
| *Nettapus auritus* | GO:0050661 | 0.02597403 | MF | NADP binding |
| *Nettapus auritus* | GO:0004812 | 0.02897103 | MF | aminoacyl-tRNA ligase activity |
| *Nettapus auritus* | GO:0016875 | 0.02897103 | MF | ligase activity, forming carbon-oxygen bonds |
| *Nettapus auritus* | GO:0047485 | 0.02897103 | MF | protein N-terminus binding |
| *Nettapus auritus* | GO:0005539 | 0.02997003 | MF | glycosaminoglycan binding |
| *Nettapus auritus* | GO:0004857 | 0.03096903 | MF | enzyme inhibitor activity |
| *Nettapus auritus* | GO:0008081 | 0.03096903 | MF | phosphoric diester hydrolase activity |
| *Nettapus auritus* | GO:0015631 | 0.03096903 | MF | tubulin binding |
| *Nettapus auritus* | GO:0035198 | 0.03096903 | MF | miRNA binding |
| *Nettapus auritus* | GO:0008373 | 0.03196803 | MF | sialyltransferase activity |
| *Nettapus auritus* | GO:0003887 | 0.03296703 | MF | DNA-directed DNA polymerase activity |
| *Nettapus auritus* | GO:0005179 | 0.03296703 | MF | hormone activity |
| *Nettapus auritus* | GO:0019239 | 0.03596404 | MF | deaminase activity |
| *Nettapus auritus* | GO:0052689 | 0.03596404 | MF | carboxylic ester hydrolase activity |
| *Nettapus auritus* | GO:0008509 | 0.03696304 | MF | anion transmembrane transporter activity |
| *Nettapus auritus* | GO:0051787 | 0.03696304 | MF | misfolded protein binding |
| *Nettapus auritus* | GO:0017124 | 0.03796204 | MF | SH3 domain binding |
| *Nettapus auritus* | GO:0051219 | 0.03896104 | MF | phosphoprotein binding |
| *Nettapus auritus* | GO:0060590 | 0.03896104 | MF | ATPase regulator activity |
| *Nettapus auritus* | GO:0140101 | 0.03896104 | MF | catalytic activity, acting on a tRNA |
| *Nettapus auritus* | GO:0000049 | 0.03996004 | MF | tRNA binding |
| *Nettapus auritus* | GO:0005342 | 0.03996004 | MF | organic acid transmembrane transporter activity |
| *Nettapus auritus* | GO:0046943 | 0.03996004 | MF | carboxylic acid transmembrane transporter activity |
| *Nettapus auritus* | GO:0001221 | 0.04095904 | MF | transcription cofactor binding |
| *Nettapus auritus* | GO:0008201 | 0.04095904 | MF | heparin binding |
| *Nettapus auritus* | GO:0016616 | 0.04095904 | MF | oxidoreductase activity, acting on the CH-OH group of donors, NAD or NADP as acceptor |
| *Nettapus auritus* | GO:0048306 | 0.04095904 | MF | calcium-dependent protein binding |
| *Nettapus auritus* | GO:0003684 | 0.04195804 | MF | damaged DNA binding |
| *Nettapus auritus* | GO:0003743 | 0.04195804 | MF | translation initiation factor activity |
| *Nettapus auritus* | GO:0015081 | 0.04195804 | MF | sodium ion transmembrane transporter activity |
| *Nettapus auritus* | GO:0016614 | 0.04195804 | MF | oxidoreductase activity, acting on CH-OH group of donors |
| *Nettapus auritus* | GO:0036002 | 0.04195804 | MF | pre-mRNA binding |
| *Nettapus auritus* | GO:0043425 | 0.04195804 | MF | bHLH transcription factor binding |
| *Nettapus auritus* | GO:0009881 | 0.04295704 | MF | photoreceptor activity |
| *Nettapus auritus* | GO:0016829 | 0.04295704 | MF | lyase activity |
| *Nettapus auritus* | GO:0020037 | 0.04295704 | MF | heme binding |
| *Nettapus auritus* | GO:0034061 | 0.04295704 | MF | DNA polymerase activity |
| *Nettapus auritus* | GO:0070888 | 0.04295704 | MF | E-box binding |
| *Nettapus auritus* | GO:0004842 | 0.04395604 | MF | ubiquitin-protein transferase activity |
| *Nettapus auritus* | GO:0015662 | 0.04395604 | MF | ion transmembrane transporter activity, phosphorylative mechanism |
| *Nettapus auritus* | GO:0016298 | 0.04395604 | MF | lipase activity |
| *Nettapus auritus* | GO:0046906 | 0.04395604 | MF | tetrapyrrole binding |
| *Nettapus auritus* | GO:0015085 | 0.04495505 | MF | calcium ion transmembrane transporter activity |
| *Nettapus auritus* | GO:0019787 | 0.04495505 | MF | ubiquitin-like protein transferase activity |
| *Nettapus auritus* | GO:0004386 | 0.04595405 | MF | helicase activity |
| *Nettapus auritus* | GO:0004860 | 0.04595405 | MF | protein kinase inhibitor activity |
| *Nettapus auritus* | GO:0008094 | 0.04595405 | MF | DNA-dependent ATPase activity |
| *Nettapus auritus* | GO:0016776 | 0.04595405 | MF | phosphotransferase activity, phosphate group as acceptor |
| *Nettapus auritus* | GO:0016830 | 0.04595405 | MF | carbon-carbon lyase activity |
| *Nettapus auritus* | GO:0019210 | 0.04595405 | MF | kinase inhibitor activity |
| *Nettapus auritus* | GO:0018024 | 0.04695305 | MF | histone-lysine N-methyltransferase activity |
| *Nettapus auritus* | GO:0032182 | 0.04695305 | MF | ubiquitin-like protein binding |
| *Nettapus auritus* | GO:0034713 | 0.04695305 | MF | type I transforming growth factor beta receptor binding |
| *Nettapus auritus* | GO:0035257 | 0.04695305 | MF | nuclear hormone receptor binding |
| *Nettapus auritus* | GO:0042054 | 0.04695305 | MF | histone methyltransferase activity |
| *Nettapus auritus* | GO:0051393 | 0.04695305 | MF | alpha-actinin binding |
| *Nettapus auritus* | GO:0000979 | 0.04795205 | MF | RNA polymerase II core promoter sequence-specific DNA binding |
| *Nettapus auritus* | GO:0008017 | 0.04795205 | MF | microtubule binding |
| *Nettapus auritus* | GO:0008135 | 0.04795205 | MF | translation factor activity, RNA binding |
| *Nettapus auritus* | GO:0008194 | 0.04795205 | MF | UDP-glycosyltransferase activity |
| *Nettapus auritus* | GO:0008276 | 0.04795205 | MF | protein methyltransferase activity |
| *Nettapus auritus* | GO:0016538 | 0.04795205 | MF | cyclin-dependent protein serine/threonine kinase regulator activity |
| *Nettapus auritus* | GO:0051087 | 0.04795205 | MF | chaperone binding |
| *Nettapus auritus* | GO:0003714 | 0.04895105 | MF | transcription corepressor activity |
| *Nettapus auritus* | GO:0004497 | 0.04895105 | MF | monooxygenase activity |
| *Nettapus auritus* | GO:0019829 | 0.04895105 | MF | ATPase-coupled cation transmembrane transporter activity |
| *Nettapus auritus* | GO:0042625 | 0.04895105 | MF | ATPase-coupled ion transmembrane transporter activity |
| *Nettapus auritus* | GO:0042805 | 0.04895105 | MF | actinin binding |
| *Nettapus auritus* | GO:0043621 | 0.04895105 | MF | protein self-association |
| *Nettapus auritus* | GO:0004175 | 0.04995005 | MF | endopeptidase activity |
| *Nettapus auritus* | GO:0016278 | 0.04995005 | MF | lysine N-methyltransferase activity |
| *Nettapus auritus* | GO:0016279 | 0.04995005 | MF | protein-lysine N-methyltransferase activity |
| *Nettapus auritus* | GO:0016407 | 0.04995005 | MF | acetyltransferase activity |
| *Nettapus auritus* | GO:0051427 | 0.04995005 | MF | hormone receptor binding |
